# Bacterial lipid effectors subvert host autophagic flux by inducing lysosomal dysfunction

**DOI:** 10.64898/2026.09.18.752599

**Authors:** Camille Pin, Christelle Marrauld, Priscilla Branchu, Chrystelle Bonnart, Corinne Rolland, Lhorane Lobjois, Simon Lachambre, Frédéric Taieb, Eric Oswald, Laure David

**Affiliations:** Université de Toulouse, INSERM, INRAE, ENVT, IRSD, Toulouse, France; Université de Toulouse, INSERM, CNRS, Infinity – Toulouse Institute for Infectious and Inflammatory Diseases, Toulouse, France; Université de Toulouse, CHU Toulouse, INSERM, INRAE, ENVT, IRSD, Toulouse, France

**Keywords:** Autophagy, Gram-negative bacteria, lysosomal defects, lysosomal membrane permeabilization, outer-membrane vesicle

## Abstract

HlyF and CprA, virulence factors from *Escherichia coli* and *Pseudomonas aeruginosa* respectively, constitute a previously unrecognized family of cytoplasmic enzymes that drive virulence through the production of outer membrane vesicles (OMVs) enriched in toxic bioactive lipids. Here, we show that OMVs of bacteria producing HlyF/CprA impair host autophagic flux by targeting lysosomal function. Specifically, these OMV-associated lipids disrupt autophagosome–lysosome fusion and induce hallmark features of lysosomal dysfunctions, including defective acidification, cholesterol accumulation, and membrane permeabilization. These lipid-mediated alterations compromise lysosomal integrity and block autophagic degradation. Our findings reveal a novel mechanism by which bacterial lipids subvert lysosomal functions and host clearance pathways. Given the widespread expression of HlyF/CprA homologs across diverse pathogens, this work highlights lipid effectors as key determinants of pathogenesis.

## Introduction

Beyond its fundamental role in cellular homeostasis (1,2), macroautophagy, hereafter referred to as autophagy, serves also as a key defense mechanism against microbial pathogens through a process known as xenophagy to degrade pathogens (3–5). However, pathogens have evolved strategies to evade or even exploit autophagy, such as inhibiting phagophore formation or preventing autophagosome-lysosome fusion to avoid degradation and establish a replicative niche (3,5,6). The mechanisms generally involve the recruitment or production of proteins that enable bacteria to avoid recognition by autophagy proteins or autophagy initiation (7–10). We identified a previously unrecognized family of cytoplasmic bacterial short-chain dehydrogenases/reductases (SDRs) that drive outer membrane vesicle (OMV) biogenesis in pathogenic Gram-negative bacteria (11–13). We showed that even though the SDR enzymes HlyF and CprA are strictly cytoplasmic and thus are not found in OMVs, they act as virulence factors by promoting the production of a distinct subset of OMVs (11, 13). Moreover, we demonstrated that OMVs produced by HlyF-producing *Escherichia coli* (*E. coli*) strains (called hereafter OMVs-HlyF) inhibit autophagic flux by blocking the fusion of autophagosomes with lysosomes, thereby exacerbating inflammasome activation (12). Building on these findings, we showed more recently that this mechanism is also conserved in other pathogenic bacteria producing HlyF orthologues, such as *Pseudomonas aeruginosa* through the expression of CprA, revealing a conserved family of bacterial virulence determinants capable of disrupting autophagic flux in host cells. The specific ability of these OMVs to inhibit autophagy depends on the presence of bioactive lipids, whose synthesis is catalyzed by HlyF and CprA (12,13).

In this study, we investigated the mechanisms by which OMVs and lipids from HlyF/CprA- producing bacteria (hereafter called HlyF-lipids / CprA-lipids) inhibit the fusion of autophagosomes with lysosomes. We show that treating host cells with either these specific OMVs or lipids extracted from HlyF/CprA-producing bacterial membranes leads to a rapid accumulation of phenotypically abnormal lysosomes. Specifically, these OMV-associated bioactive lipids profoundly modulate lysosomal physiology by impairing acidification and altering membrane lipidic composition and integrity, which ultimately disrupts autophagic flux. These findings establish bacterial lipid effectors as key virulence determinants, highlighting their underappreciated role in host–pathogen interactions.

## Results

### OMVs and lipids from HlyF-producing bacteria induce LC3-II accumulation

We previously demonstrated that OMVs produced by bacteria producing HlyF or its ortholog CprA inhibit autophagic flux in host cells by blocking autophagosome–lysosome fusion (11–13). To gain further mechanistic insight into this process, we first characterized the kinetics of autophagy inhibition by monitoring LC3-II accumulation, a hallmark of autophagosome formation and impaired autophagic flux. OMVs-HlyF triggered a rapid increase in LC3-II levels, detectable within 30 minutes after treatment and reaching a maximum at 1.5 hours (Fig. 1A, Fig. 1B). While our previous work established that the autophagy-disrupting activity of OMVs from CprA-producing bacteria resides within their lipid fraction (13), it remained unclear whether these bioactive lipids require vesicular delivery or whether lipids derived directly from bacterial membranes were intrinsically active. To address this question, we extracted total lipids directly from HlyF-producing bacteria and exposed host cells to this lipid fraction. Remarkably, bacterial lipids alone reproduced the effect of OMVs-HlyF, inducing a similar time-dependent accumulation of LC3-II (Fig. 1C, Fig. 1D). These findings identify bacterial lipids as the primary bioactive effectors responsible for autophagy inhibition and demonstrate that their activity is independent of OMVs-associated delivery.

**Figure 1:**
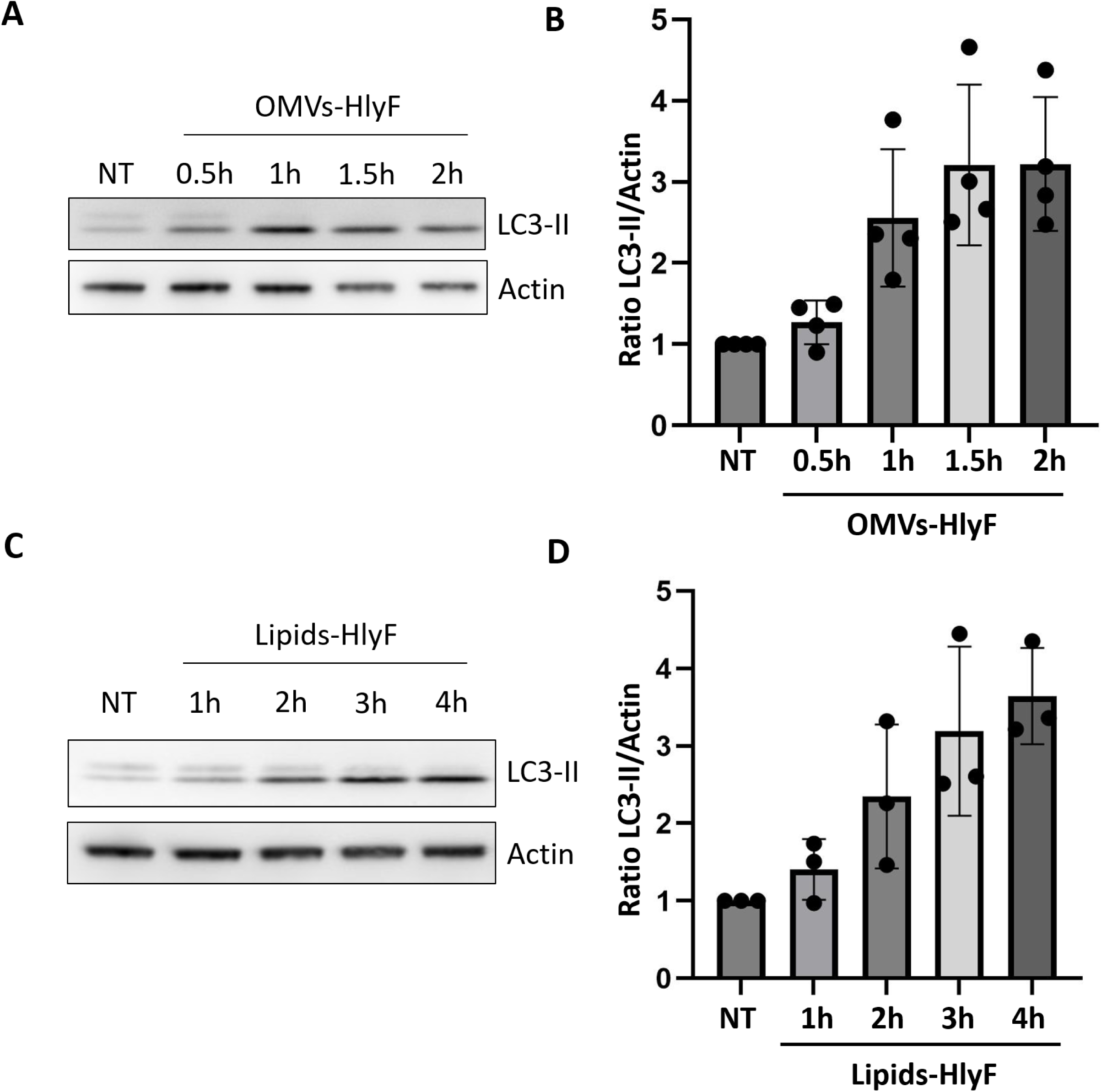
OMVs and lipids from HlyF-producing bacteria induce LC3-II accumulation. **(A)** LC3-II and actin western-blot analysis of LAMP1-GFP HeLa cells treated with OMVs-HlyF at 10 µg/mL of lipids or untreated for the times indicated. (**B)** Quantification of LC3-II/Actin of 4 independent experiments. Each dot represents one experiment. The graph shows the mean and the standard deviation for each condition. (**C)** LC3-II and actin western-blot analysis of LAMP1-GFP HeLa cells treated with lipids-HlyF at 10 µg/mL or untreated for the times indicated. (**D)** Quantification of LC3-II/Actin of 3 independent experiments. Each dot represents one experiment. The graph shows the mean and the standard deviation for each condition.

### OMVs and lipids from HlyF-producing bacteria induce lysosomes swelling

To investigate the mechanism by which OMVs and lipids from HlyF-producing *E. coli* impair the fusion between autophagosomes and lysosomes (12,13), we examined their effect on lysosomal morphology in host cells. We used HeLa cells stably producing LAMP1-GFP, which enables direct visualization of the green fluorescence associated with the late endosome and lysosome membrane protein LAMP1. Treatment for 1 hour with OMVs-HlyF and 3 hours with lipids-HlyF resulted in the formation of 30% larger LAMP1-GFP-positive foci compared to untreated cells (Fig. 2A, 2B).

**Figure 2:**
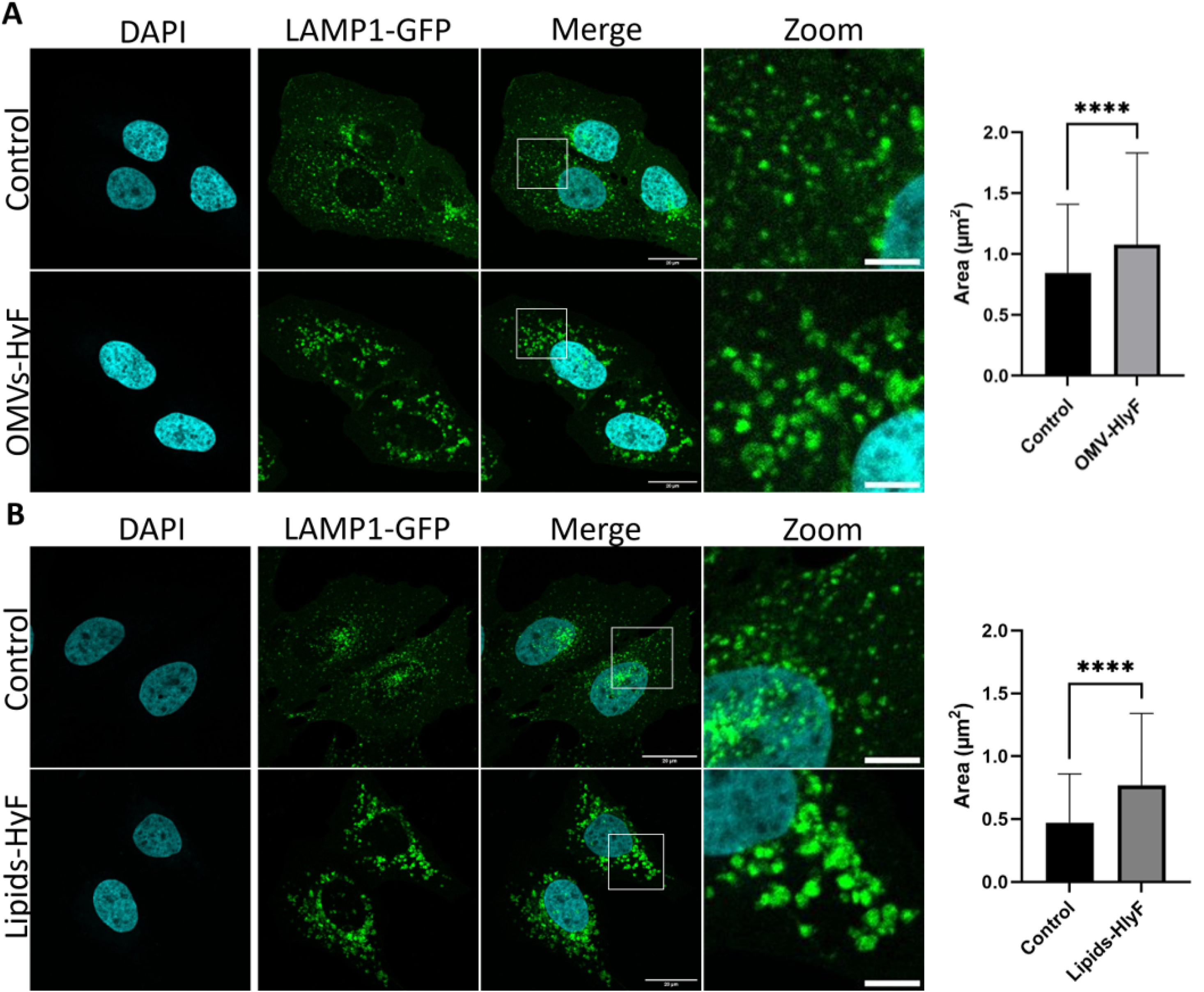
OMVs and lipids from HlyF-producing bacteria induce lysosomes swelling. **(A)** LAMP1-GFP HeLa cells were treated with an equivalent quantity of OMVs-HlyF at 10 µg/mL lipids for 1 hour or untreated. (**B)** LAMP1-GFP HeLa cells were treated with an equivalent quantity of lipids-HlyF at 10 µg/mL for 3 hours or untreated. Pictures were acquired with Zeiss LSM 710 confocal microscope. Scale bar = 20 µm. Boxed regions are shown at higher magnification (scale bar = 5µm). Images representative of at least 3 independent experiments. The graphs show the mean and the standard deviation of lysosomes areas in each condition. **** p < 0.0001 (Welch’s t-test).

These findings support the fact that OMVs-HlyF contain specific biolipids that are responsible for the enlargement of LAMP1 positive vesicles.

### OMVs and lipids from HlyF-producing bacteria induce a loss of lysosomes acidification

Enlargement of lysosomes is often associated with lysosomal dysfunctions (19–23). We next investigated lysosomal functions in cells treated with OMVs-HlyF and lipids-HlyF. To assess whether lysosomes in OMVs-HlyF-treated cells maintained an acidic content, we used LysoTracker, a pH-sensitive lysosomal probe. HeLa cells producing LAMP1-GFP treated for 1 hour with OMVs-HlyF (Fig.3A) or lipids-HlyF (Fig. 3B) presented a marked decrease in the LysoTracker signal compared to untreated cells.

**Figure 3:**
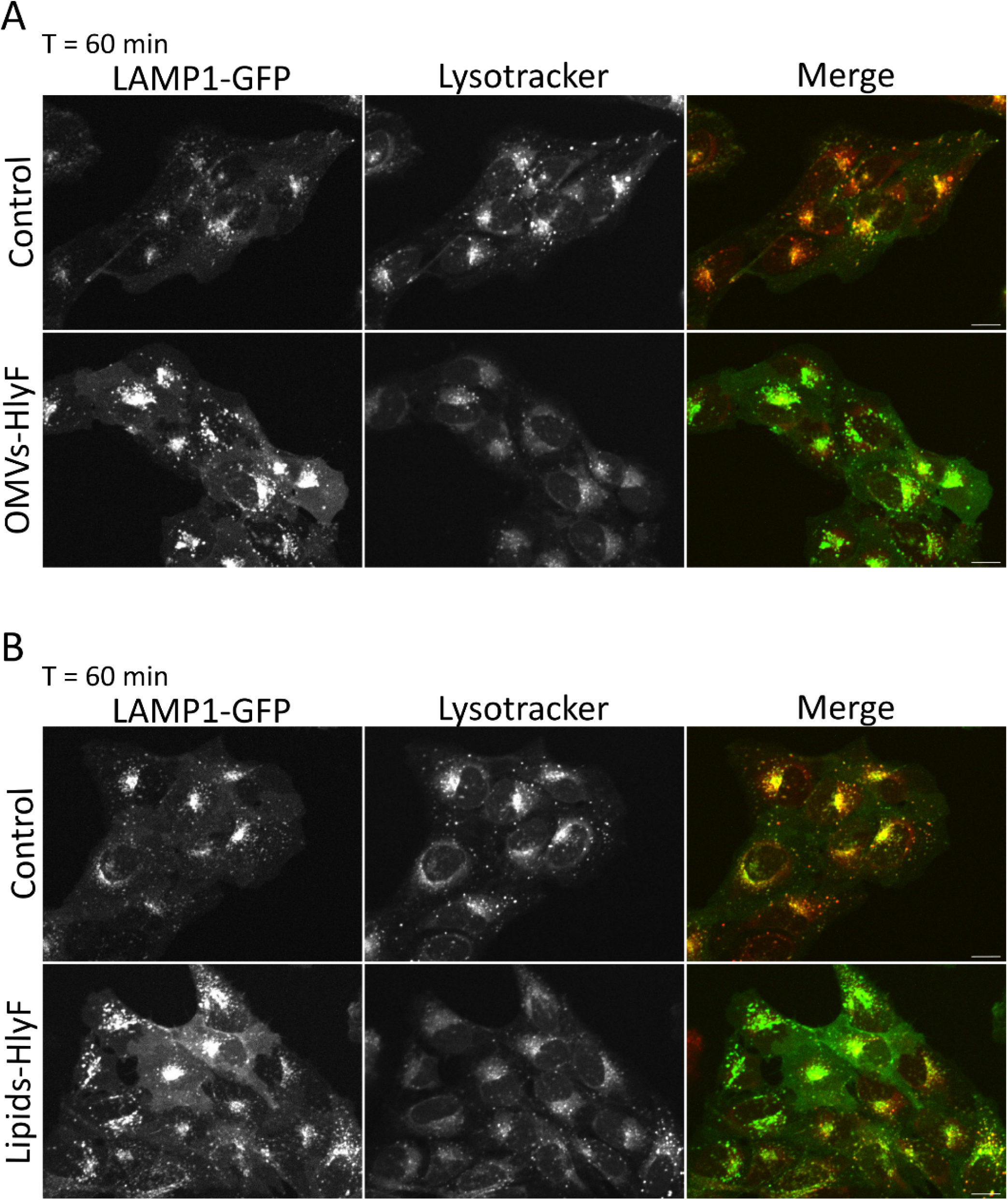
OMVs and lipids from HlyF-producing bacteria induce a loss of lysosomes acidification. **(A-B)** LAMP1-GFP HeLa cells were treated for 1 hour with **(A)** OMVs-HlyF at 10 µg/mL lipids or **(B)** lipids- HlyF at 10 µg/mL or untreated. LAMP1-GFP and LysoTracker signals are displayed in green and red respectively in the merged channel. Experiments were performed with a Spinning disk confocal set up to incubate cells at 37°C with 5% CO_2_ during the experiment. Images representative of at least 3 independent experiments. Scale bar = 16 µm.

Using live-cell imaging, we found that the LysoTracker signal declined within ∼30 minutes following OMVs-HlyF or lipids-HlyF treatment and was nearly abolished irreversibly after 1.5 hours, in contrast to untreated cells (Fig. S1). These results indicate a rapid and pronounced functional defect in lysosomes, characterized by loss of acidic pH shortly after exposure to OMVs-HlyF or their lipid components. These findings prompted us to investigate the mechanisms associated with this loss of lysosomal function.

### OMVs and lipids from HlyF-producing bacteria induce cholesterol accumulation in lysosomes

Lysosomes play a central role in cellular cholesterol homeostasis, and the accumulation of cholesterol within these organelles is a hallmark of several lysosomal storage disorders. Importantly, lysosomal cholesterol accumulation has also been associated with defective lysosomal acidification, impaired autophagic flux, and alterations in lysosomal membrane integrity (24–28). We therefore investigated whether OMVs-HlyF and their bioactive lipid components affect lysosomal cholesterol homeostasis in host cells. To visualize intracellular cholesterol distribution, free (unesterified) cholesterol was stained with filipin III. As expected, and consistent with the inability of bacteria to synthesize cholesterol, filipin III did not label HlyF-associated bacterial lipids (Fig. S2). Following a 1-hour treatment with OMVs- HlyF, we observed a marked redistribution of cholesterol compared to untreated cells (Fig. 4A). Whereas control cells displayed diffuse filipin III staining with limited perinuclear signal, OMVs-HlyF-treated cells exhibited a pronounced accumulation of bright filipin III– positive foci in the perinuclear region. Furthermore, filipin III staining colocalized with LAMP1-GFP fluorescence in treated cells (Fig. 4A), indicating a marked accumulation of cholesterol within lysosomes. Consistently, in LAMP1-GFP HeLa cells, lipids-HlyF treatment altered intracellular cholesterol distribution, leading to colocalization of filipin III and LAMP1-GFP signals, in contrast to untreated cells (Fig. 4B).

**Figure 4:**
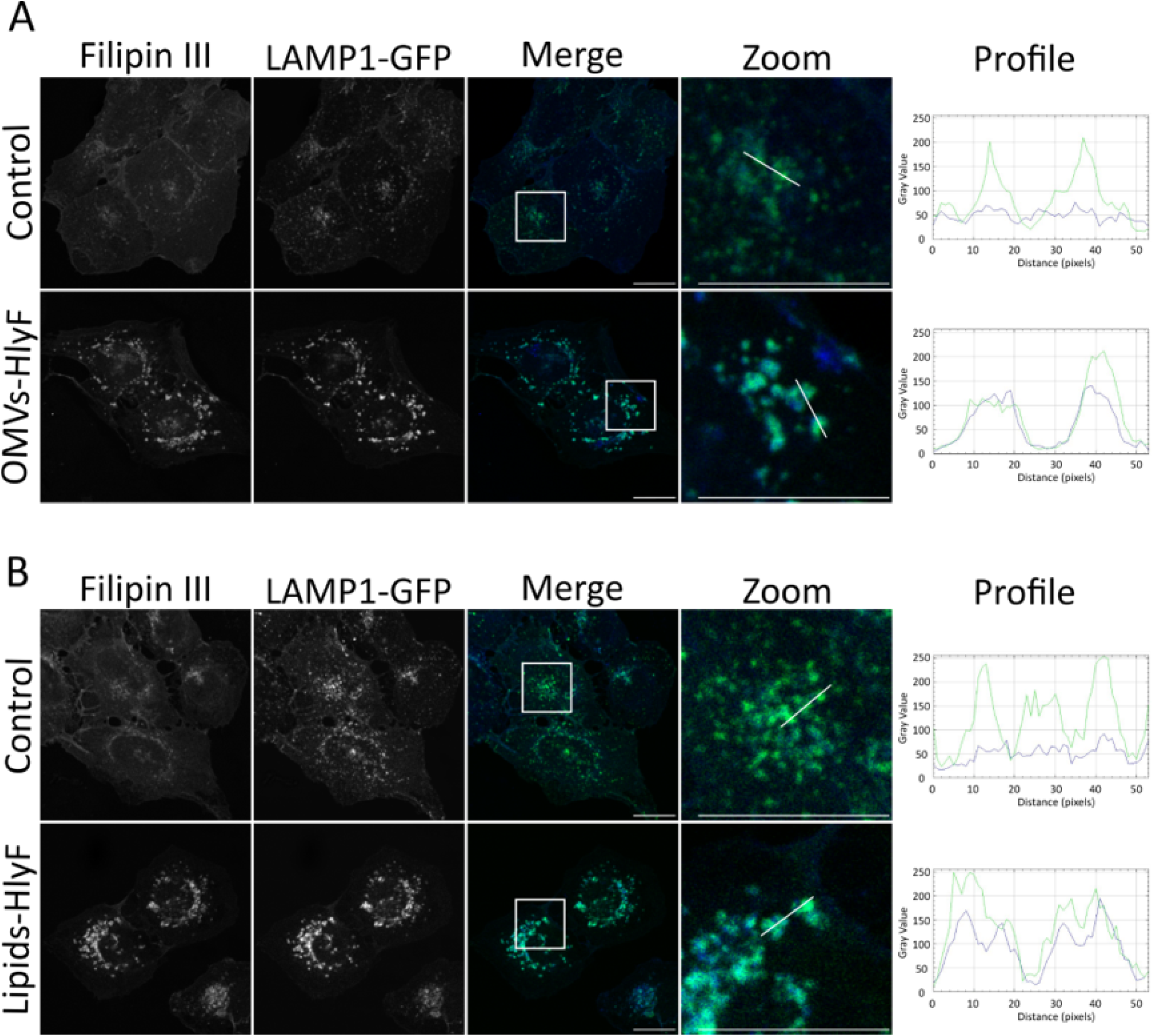
OMVs and lipids from HlyF-producing bacteria induce cholesterol accumulation in lysosomes. **(A-B)** LAMP1-GFP HeLa cells were treated for 1 hour with **(A)** OMVs-HlyF at 10 µg/mL lipids or **(B)** lipids- HlyF at 10 µg/mL or untreated. Cells were then subsequently fixed and labeled with 50 µg/mL filipin III. Images are representative of three independent experiments. Boxed regions are shown at higher magnification. Filipin III and LAMP1-GFP signals are displayed in blue and green respectively in the merged channel and profiles sections. Profile color histograms were generated from the lines shown in Zoom using ImageJ RGB Profiler plugin. The x-axis corresponds to the number of pixels in the slice, equivalent to 7 µm, and the y-axis represents the grayscale intensity. Scale bar = 20 µm.

These findings demonstrate that HlyF-dependent bacterial lipids are sufficient to disrupt lysosomal cholesterol homeostasis independently of OMV-associated delivery, further identifying these lipids as the primary bioactive effectors underlying the lysosomal alterations induced by HlyF-producing bacteria. Because lysosomal cholesterol accumulation is increasingly recognized as an early hallmark of lysosomal membrane dysfunction preceding membrane permeabilization, we therefore next investigated whether these bacterial lipids induce lysosomal membrane permeabilization, thereby initiating the cascade of events leading to impaired lysosomal function, notably loss of acidification, and autophagy blockade (23–28).

### OMVs or lipids from HlyF-producing bacteria induce lysosomal membrane permeabilization

To assess lysosomal membrane integrity, we monitored the recruitment of galectins, a family of β-galactoside-binding lectins that translocate to damaged lysosomes by binding to luminal glycans exposed upon membrane rupture (29,30). We studied specifically galectin-1 and galectin-3 recruitment to lysosomes. Under homeostatic conditions, galectin-1 and galectin-3 display diffuse cytoplasmic distributions; however, lysosomal membrane permeabilization (LMP) triggers their rapid coalescence into discrete puncta at sites of disruption (Fig. 5A). To determine whether OMVs-HlyF and lipids-HlyF compromise lysosomal integrity, we analyzed galectin-1 localization following respectively 1.5 and 4 hours of treatment. In treated cells, galectin-1 formed prominent cytoplasmic foci (absent in untreated controls) that colocalized with LAMP1-GFP puncta, confirming their lysosomal identity (Fig. 5B and Fig. 5C). Strikingly, while galectin-1 and galectin-3 both serve as LMP sensors, they differ in their binding properties and sensitivity: galectin-1 is a homodimeric galectin with a single carbohydrate recognition domain (CRD), whereas galectin-3 is an only chimera-type galectin, composed of a single CRD fused to an N-terminal oligomerization domain (30). Consistent with this, galectin-3 lysosomal recruitment pattern was different from the one associated to galectin-1 recruitment in cells exposed to OMVs-HlyF (Fig. 3), suggesting that OMVs-HlyF induce a form of lysosomal membrane damage that differentially recruits these two galectins.

**Figure 5:**
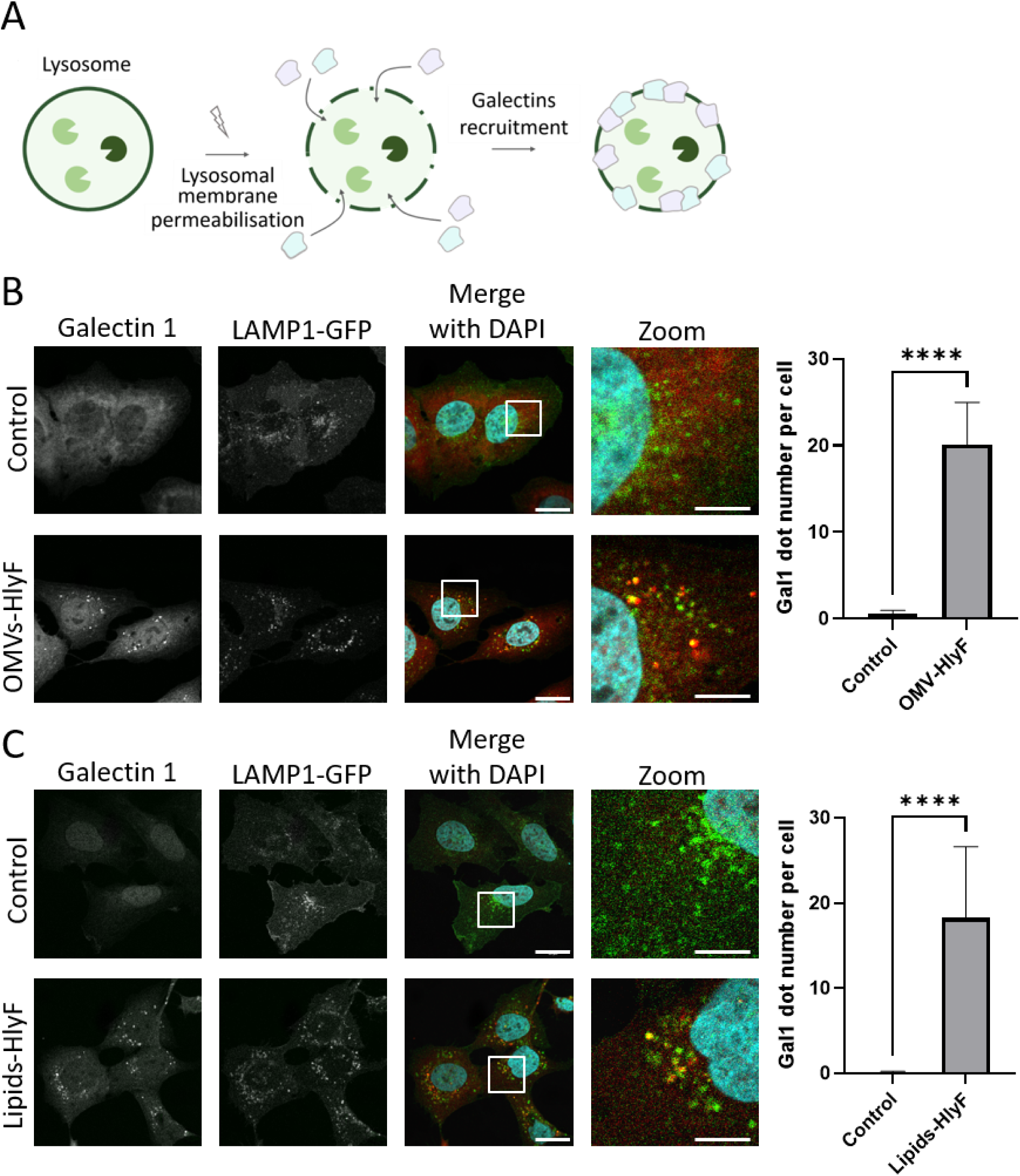
OMVs or lipids from HlyF-producing bacteria induce lysosomal membrane permeabilization. **A**. Model of galectin recruitment: Upon lysosomal membrane permeabilization, cytosolic galectins, represented in green and pink, are recruited to fill the gaps in the lysosomal membrane. This recruitment leads to the concentration of galectins associated with lysosomes, as evidenced by the appearance of bright galectin foci. **B-C**. LAMP1-GFP HeLa cells were treated for (**B**) 1.5 hours with OMVs-HlyF at 10 µg lipids /mL or untreated or (**C**) for 4 hours with lipids-HlyF at 10 µg/mL or untreated. Cells were then fixed and labeled with an anti- Galectin-1 antibody. Galectin-1 and LAMP1-GFP are displayed in red and green respectively in the merged channel. Scale bar = 20 µm. Boxed regions are shown at higher magnification (scale bar = 5µm). Images are representative of three independent experiments. Scale bar = 20 µm. The graphs show the mean and the standard deviation of the number of galectin-1 dots per cell in each condition. **** p < 0.0001 (Welch’s t-test).

### Lysosomal membrane damage observed with by HlyF-derived bacterial lipids is reproduced by CprA-derived bacterial lipids and conserved across eukaryotic cell types

We previously established that CprA is a functional ortholog of HlyF that promotes the production of OMVs with the ability to block autophagic flux (13). To determine whether this functional conservation reflects a shared capacity to generate bioactive lipids targeting host lysosomal function, we isolated lipid extracts from CprA-producing bacteria (lipids-CprA) and assessed their activity in HeLa cells. Strikingly, lipids-CprA reproduced the phenotype induced by lipids-HlyF, triggering rapid recruitment of galectin-1 to lysosomal membranes (Fig. 6A), a hallmark of lysosomal membrane permeabilization (LMP). These results demonstrate that the ability to generate lysosome-damaging bacterial lipids is an evolutionarily conserved property of the HlyF/CprA ortholog family, indicating that lipid remodeling represents a common virulence mechanism associated with these functionally related bacterial virulence factors. We next investigated whether this lysosomal targeting activity was restricted to epithelial cells or represented a broader host response. OMVs produced by HlyF-producing bacteria induced lysosomal membrane destabilization and galectin-1 recruitment not only in epithelial cells but also in immune cells, including THP-1- derived macrophages (Fig. 6B) and in primary cells such as the mouse dorsal root ganglion (DRG) derived cells, containing neurons and glial cells (Fig. 6C). Together, these findings demonstrate that HlyF-associated bacterial lipids target a conserved lysosomal vulnerability across diverse mammalian cell types.

**Figure 6:**
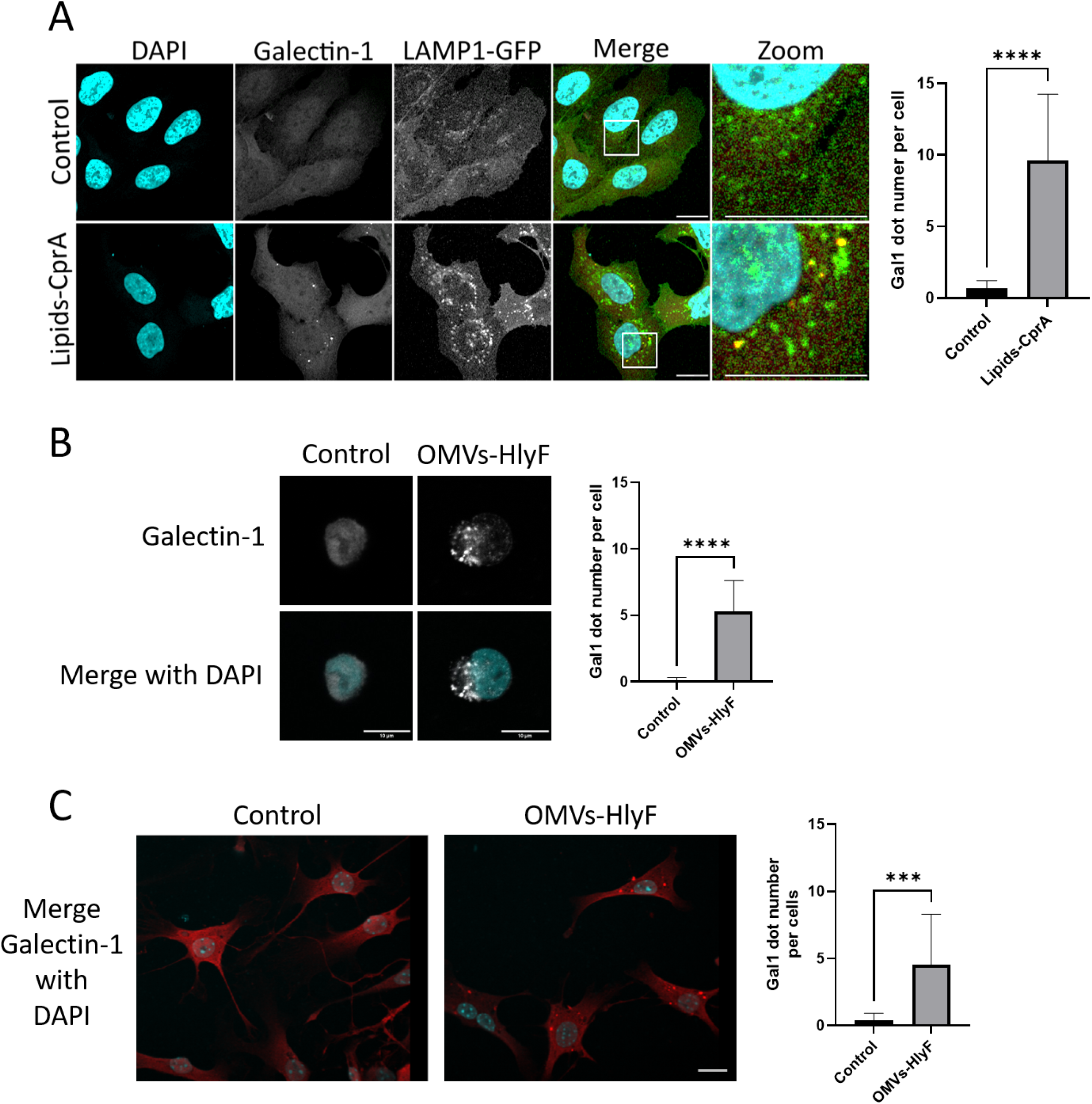
Lysosomal membrane damage observed with by HlyF-derived bacterial lipids is reproduced by CprA-derived bacterial lipids and conserved across eukaryotic cell types. (A) LAMP1-GFP HeLa cells were exposed to 10µg/mL of Lipids-CprA for 1.5 hours before being fixed and labeled with an anti-Galectin-1 antibody. Galectin-1 and LAMP1-GFP are displayed in red and green respectively in the merged channel. Cell nuclei were stained with DAPI (cyan). Images are representative of three independent experiments. Boxed regions are shown at higher magnification. Scale bar = 20 µm. The graph shows the mean and the standard deviation of the number of Galectin-1 dots per cell in each condition. **** p < 0.0001 (Welch’s t-test). (B) THP1 cells were exposed to 20µg/mL OMVs-HlyF for 3hours before being fixed and labeled with an anti- Galectin-1 antibody (grey signal). Cell nuclei were stained with DAPI (cyan). Images are representative of two independent experiments. Scale bar = 10 µm. The graph shows the mean and the standard deviation of the number of Galectin-1 dots per cell in each condition. **** p < 0.0001 (Welch’s t-test). (C) Murine DRG cultures were infected with 10µg/mL OMVs-HlyF for 3 hours. Cells were fixed in PFA and immunostained with anti- Galectin-1 antibodie (red signal). Cell nuclei were stained with DAPI (cyan). The graph is representative of two independent experiments. ***p<0.001 (Kolmogorov-Smirnov test).

### HlyF-derived lipids integrate into lysosomal membranes and alter their structural organization

To elucidate the mechanism by which HlyF-derived bioactive lipids damage lysosomal membranes, we investigated whether they were directly incorporated into the lysosomal membrane bilayer. To track their intracellular fate, we labeled the lipid extracts with DiI, a hydrophobic fluorescent dye that intercalates stably into lipid bilayers. The lipid mixture were extruded to generate uniform unilamellar liposomes of approximately 100 nm in diameter (Fig. 7A). These liposomes (liposomes-HlyF) were formulated with 50% HlyF-derived bacterial lipids and 50% of synthetic lipids representing the major components of the outer membrane of *E. coli* (60% POPE, 30% POPG, 10% cardiolipin), as described by Kehl et al. (18). Control liposomes (liposomes-control) were generated from 100% synthetic lipids of identical composition, serving as a baseline for membrane incorporation behavior. Upon uptake by host cells, DiI fluorescence was detected in association with lysosomes under both conditions, but with strikingly distinct distribution patterns that reveal fundamentally different fates. In liposomes-control-treated cells, DiI signal was confined to the lysosomal lumen, consistent with delivery of vesicles to lysosomes by endosomal engulfment (Fig. 7B–C). In contrast, liposomes-HlyF-treated cells displayed DiI-positive arcs specifically localized within the lysosomal limiting membrane, concentrated in regions of reduced LAMP1-GFP signal, a pattern indicative of membrane intercalation rather than luminal deposition (Fig. 7D–E). This finding was independently validated using a fluorescently labeled synthetic lipid incorporated directly into the liposome formulation, yielding equivalent results (Fig. 4). Collectively, these data demonstrate that HlyF-derived lipids possessed an intrinsic capacity to integrate into lysosomal membranes, where they induced structural perturbations that likely underly the lysosomal cholesterol accumulation, membrane permeabilization, and autophagic blockade described above.

**Figure 7:**
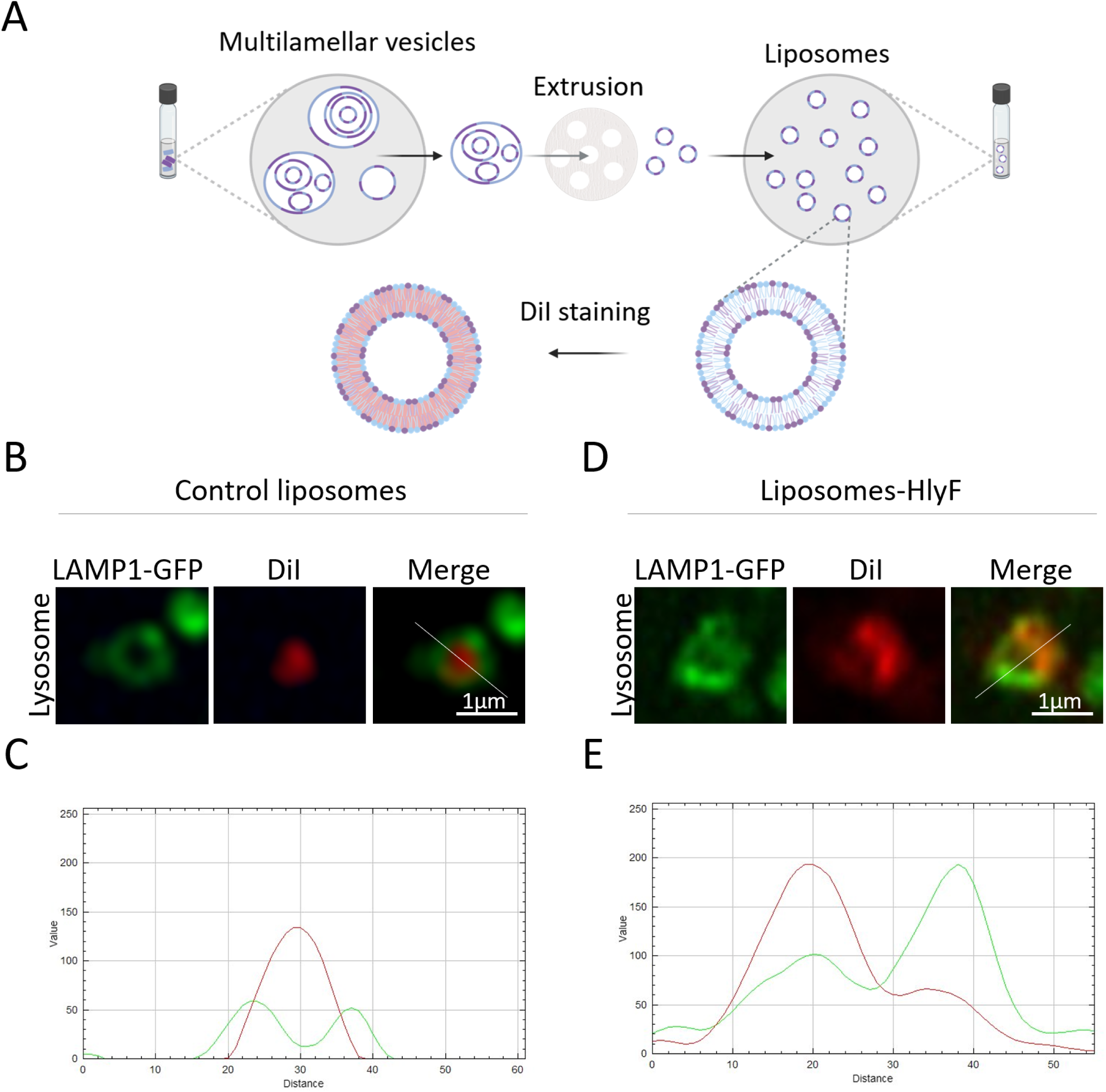
HlyF-derived lipids integrate into lysosomal membranes and alter their structural organization. **(A)**. To prepare liposomes, lipids were resuspended in aqueous media and then extruded though 200 then 100 nm filters. This step allows to break the multilamellar vesicles produced during the resuspension of lipids. After extrusion, liposomes present a homogeneous size. Liposomes were then labelled with the hydrophilic probe DiI to stain their lipids bilayer. Created with BioRender.com. (**B-E**) LAMP1-GFP HeLa were treated with (**B)** 100 µg/mL DiI-stained liposomes made with 100% of synthetic lipids (Control liposomes) or (**D)** 50% of synthetic lipids added to 50% of lipids-HlyF (liposomes-HlyF) during 3 hours. Images were acquired with a Zeiss 880 confocal microscope with an Airyscan module. Scale bar = 1 µm. (**C, E**) Profiles were made with the RGB profiler plugin in ImageJ and represent LAMP1-GFP and DiI signals in green and red respectively. Distance are expressed in pixels.

## Discussion

In this study, we sought to elucidate the mechanism by which bacteria producing HlyF or CprA interfere with the host autophagic flux (11–13). We demonstrated that both OMVs and lipid extracts from these bacteria induced autophagosome accumulation and displayed multiple hallmarks of lysosomal dysfunction. Specifically, exposure to OMVs-HlyF or lipids- HlyF resulted in lysosomal enlargement, defective acidification, cholesterol accumulation, and ultimately lysosomal membrane permeabilization.

Our findings identify HlyF- and CprA-dependent bacterial lipids as key effectors responsible for lysosomal dysfunction. Rather than acting through a conventional receptor-mediated mechanism, these bacterial lipids appear to directly alter the physical properties of the lysosomal limiting membrane itself. We propose that their incorporation into lysosomal membranes affects membrane organization, thereby affecting the biophysical properties required for proper lysosomal function. This model is consistent with the emerging concept that membrane lipids are not passive structural components but active regulators of organelle identity, membrane dynamics, and signaling pathways, particularly within the endolysosomal and autophagic systems (31–33). The consequences of this membrane remodeling are likely multifactorial and interconnected. Alteration of the lysosomal lipid environment may affect the organization and activity of membrane-associated protein complexes required for lysosomal maturation, autophagosome–lysosome fusion, and maintenance of lysosomal homeostasis. In addition, cholesterol accumulation represents a critical amplifier of lysosomal dysfunction, as excessive lysosomal cholesterol has been linked to impaired lysosomal motility, altered motor protein recruitment, defective SNARE organization, and disrupted pH regulation (19,21,22,29,34). These alterations may progressively compromise lysosomal function. Consistent with this interpretation, the redistribution of LAMP1 observed after exposure to HlyF-dependent bacterial lipids suggests a broader remodeling of lysosomal membrane organization, potentially affecting the localization and activity of resident lysosomal proteins (26,27,30). An important question is how bacterial lipids access the lysosomal limiting membrane. Lysosomal membranes undergo continuous renewal through membrane trafficking pathways involving plasma membrane-derived endosomes, recycling compartments, and endosomal maturation. We propose that exogenously delivered bacterial lipids can exploit these physiological trafficking routes to reach and accumulate within lysosomal membranes without requiring bacterial invasion (Fig. 8). Collectively, our data support a model in which HlyF- and CprA-family virulence factors promote the production of bioactive bacterial lipids that remodel host lysosomal membranes, leading to membrane destabilization, loss of lysosomal function, and blockade of autophagic flux. This mechanism reveals lipid remodeling as a conserved bacterial strategy to target a fundamental cellular organelle and highlights the lysosomal membrane as a previously underappreciated target of bacterial virulence factors.

**Figure 8:**
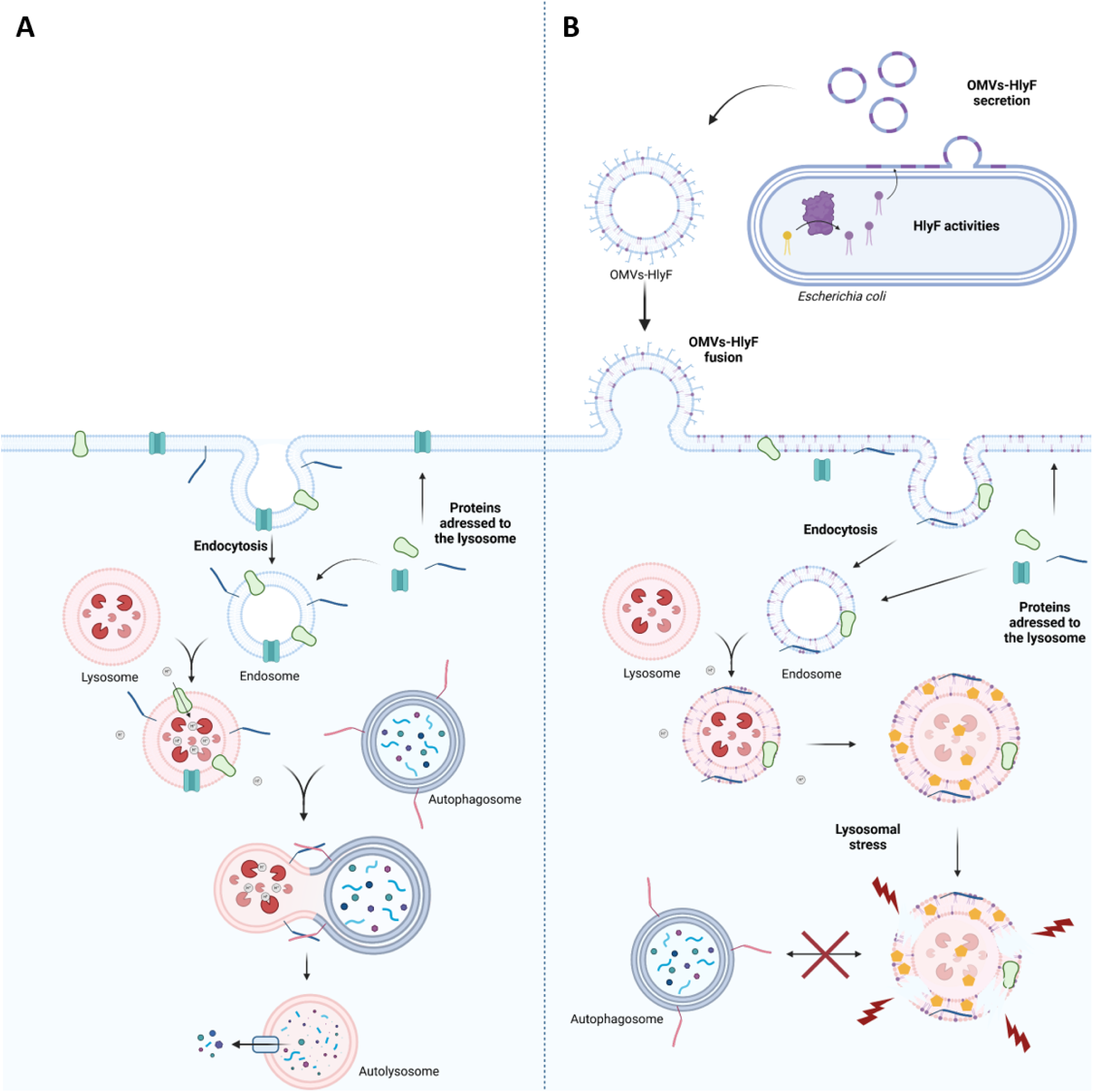
Model of the mode of action of OMVs from HlyF/CprA-producing bacteria on lysosomes leading to autophagic blockage. **(A)** In control cells, lysosome maturation is dependent of endocytosis. A subset of proteins and membranes destined for lysosomes are first trafficked to the plasma membrane before being internalized and delivered to endosomes. Then, endosomes fuse with lysosomes allowing lysosomal maturation and functionality. (**B)** In bacteria producing HlyF, this enzyme may generate a lipid pool (lipids-HlyF), distinct from lipids produced by a bacteria that does not produce HlyF. This lipid pool is subsequently incorporated into the bacterial outer membrane. Upon production, outer membrane vesicles (OMVs) are enriched with these modified lipids, forming OMVs-HlyF. These OMVs-HlyF fuse with the host cell plasma membrane, delivering lipids-HlyF into the membrane. From there, lipids are internalized via endocytosis and trafficked through the endosomal system. Fusion of endosomes with lysosomes delivers lipids-HlyF to lysosomal membranes, where they induce disturbance of membrane lipids and disrupt pH homeostasis. This is followed by cholesterol accumulation within lysosomes and eventual membrane permeabilization. These lipid-induced lysosomal defects ultimately prevent lysosome - autophagosome fusion, leading to a blockade of autophagic flux. Created with BioRender.com.

Previous studies have shown that bacterial strains producing HlyF or CprA are associated with severe acute infections in both humans and animals (11,13,31). In murine models of sepsis and urosepsis, deletion of *hlyF* or *cprA* significantly improved clinical outcomes and survival (13,31). Moreover, blockade of autophagy amplifies the host inflammatory response by disrupting its negative regulation of inflammasome activation, thereby likely contributing to disease progression and the onset of severe conditions such as sepsis (12,31). These findings underscore the critical role of HlyF and CprA in the virulence of Gram-negative bacteria. As autophagy is a key innate defense mechanism against pathogens, it is not surprising that pathogens have evolved strategies to subvert this process for their own benefit. It is well-established that both bacterial, parasitic and viral pathogens can inhibit autophagosome-lysosome fusion to avoid degradation and establish intracellular replication niches (31–33). Consistent with this strategy, *E. coli* strains producing HlyF exhibit improved survival capacity within macrophages (35,36), likely due to their ability to evade lysosomal degradation through autophagy inhibition. Moreover, beyond their role in autophagy blockade, our findings suggest that HlyF- and CprA-dependent bioactive lipids alter drastically lysosomal homeostasis in host cells. Given the central role of the autophagy-lysosome pathway in restraining LPS-induced inflammatory signaling, inflammasome activation, and pyroptotic cell death, such alterations are likely to impair the ability of immune cells to properly coordinate their response to PAMP detection (23,37–40). In particular, autophagy and lysosomal dysfunctions have been linked to the exacerbation of inflammasome activation through uncontrolled induction of pyroptosis, which can lead to aggravation of an infection and even sepsis (12). Collectively, these data suggest that, by destabilizing lysosomal membranes and impairing lysosomal function, HlyF- or CprA- dependent bioactive bacterial lipids may therefore not only facilitate bacterial persistence by avoiding autophago-lysosomal degradation (35,36) but also exacerbate the magnitude of pyroptotic response and inflammation activation during infection, thereby contributing to disease severity which may ultimately progress to sepsis (12).

Although bacteria able to produce these lysotoxic lipids have been primarily associated with acute infections, they are also present in the gut microbiota as asymptomatic colonizers.

Notably, analysis of a collection *E. coli* strains isolated from healthy asymptomatic human beings revealed that the gene coding for HlyF is present in the genome of ∼20% of the isolates (41). Given that defects in autophagy and lysosomal pathways are implicated in metabolic disorders such as non-alcoholic fatty liver disease, as well as neurodegenerative diseases including Parkinson’s and Alzheimer’s (19,38–41), the sustained release of these bioactive bacterial lipids may represent a previously underappreciated long-term risk. OMVs produced by Gram-negative bacteria, including *E. coli*, have been detected in distant organs such as the liver, heart, and brain (42,43). This systemic distribution raises particular concern for OMVs derived from HlyF-producing bacteria, which carry lipids capable of disrupting autophagy and lysosomal function across multiple cell types, including peripheral nervous system cells derived from mouse dorsal root ganglia, highlighting the conserved lysotoxic activity of OMVs and of bioactive lipids in primary cells. The chronic, even low-level, production and secretion of such lipids, as free molecules or via OMVs, by commensal bacteria suggests a potential mechanism for their gradual diffusion and accumulation in host tissues over time, particularly in long-lived cells, initially in cells close to the site of bacterial colonization, then potentially spreading further and further via direct spread or through the bloodstream. Together, these observations may raise important questions regarding the contribution of bacterial-derived lipids to systemic metabolic dysregulation and age-related pathologies, an area that remains largely unexplored and warrants further investigation.

In conclusion, while microorganisms are known to induce lysosomal membrane rupture (44,45), to our knowledge, this study presents the first evidence that lysosomal dysfunction can be triggered by OMVs, and more specifically, by a bacterial lipid delivered remotely from the primary site of infection. Our findings uncover a previously unrecognized mechanism by which bacterial lipids can subvert host cellular clearance pathways. This represents a novel and conserved strategy of lysosomal interference that may be employed by a wide range of pathogens. Furthermore, our work contributes to the growing recognition of bacterial extracellular vesicles as active players in chronic diseases. While the role of host-microbe interactions in chronic pathologies is increasingly acknowledged, the specific contribution of bacterial lipids, especially those with lysotoxic activity, has remained largely unexplored. The demonstration that certain bacterial OMV-associated lipids can directly impair lysosomal integrity opens new avenues for investigating the microbial etiology of metabolic and neurodegenerative disorders, and emphasizes the need to consider bacterial components in the study of non-infectious chronic diseases.

## Material and methods

### Cell culture

LAMP1-GFP HeLa cells were kindly provided by Andrea Ballabio. HeLa cells and LAMP1- GFP HeLa cells were cultured in DMEM Glutamax High glucose (Gibco 10566016) complement with 10% of fetal bovine serum (FBS) (17479633 fisher scientific) and 1% of non-essential amino-acids bovine serum (11140035 Invitrogen). One day before experiments, cells were seeded in 12-well plates with microscopy slides at 50 000 cells/mL.

THP1 cells were cultured in RPMI with Glutamax (Life Technology, 12,027,599) supplemented with 10% FBS. One day before the experiment, THP1 cells were seeded at a density of 3 × 10^5^ cells per mL.

### Dorsal root ganglion (DRG) culture and OMV treatment

Isolated DRGs were digested in L-cystein (pH 7.4; Sigma Aldrich, Saint Quentin Fallavier, France), papain (4 U/mL; Worthington) for 10 minutes at 37°C. DRG were rinsed in Leibovitz L15 Medium (Invitrogen) containing 10% of FBS, then rinsed three times in HBSS (Invitrogen). A second enzymatic dissociation was performed in 4-mg/mL dispase II (Sigma Aldrich) and 1-mg/mL collagenase type I (Sigma Aldrich) for 5 minutes at 37°C, followed by mechanical dissociation. This step was repeated until complete dissociation of the DRG up to 3 times. Finally, the cell suspension was centrifuged 600rpm (62g) for 5 minutes. The pellet was then resuspended in DMEM-glutamax containing 3% FBS and cells (corresponding to a mixture of neurons and glial cells) were plated on L-polylysin (Sigma)-coated 96-well plates and grown in Matrigel (Corning)-coated 96 well plates. The plates were incubated at 37°C under 5% CO_2_ atmosphere. After 24h, the medium was replaced by FluoroBrite™ DMEM medium (Invitrogen) containing glutamax and 3% FBS and the plate was incubated for an additional period of 16h. DRGs were then treated with OMVs for 3 hours at the indicated concentration before processing to immunofluorescence experiments.

### Bacteria strains and OMVs production

We used *E. coli* BL21 (DE3) strain transformed with either the pK184 vector encoding the 6His-*hlyF* fusion protein under *hlyF* promoter or pK184 vector encoding the 6His-*cprA^PAK^* fusion protein under *hlyF* promoter as described in Goman *et al*. (13).

Bacteria were grown in M63 medium (M1015 USbio) supplemented with 0.2 % glucose (G8769 sigma) and 50 µg/mL kanamycine sulfate (UK0015 Euromedex). After 8 hours of culture at 37°C with 240 rpm agitation, cultures were centrifuged at 6000 g for 10 minutes in order to pellet bacteria. Then supernatants were filtrated through 0.22 µM filter (431097 sigma) and ultrafiltrated through 100-kDa PES membrane cassette (147065 dutscher). The filtrate supernatants were then ultracentrifugated at 170 000 g for 3 hours to pellet OMVs. OMVs pellets were resuspended in PBS without Ca^+^ and Mg^2+^ (CS1PBS02-01 eurobio). Protein and lipid concentrations in OMVs were evaluated by doing a BCA protein assay (DCTM protein assay Biorad) and a lipid dosage with a sulfo-phospho-vanillin reaction as described in Izard *et al* (46) with Triolein as a reference (44896-U Sigma Aldrich).

### Filipin III labelling

LAMP1-GFP HeLa were seeded into 12-well plates with sterile slides in the bottom of the wells the day before the experiment. The day after, cells were treated as indicated in the legends of figures. After treatment, the cells were fixed with 4% PFA for 12 minutes at RT followed by 3 washes with PBS. The cells were then incubated with 1.5 mg/mL of glycine for 10 minutes at RT and then incubated with 50 µg/mL filipin III (F4767 sigma) at RT for an extra 2-4 hours. The slides were then mounted with Fluoroshield (F6182 sigma). Images were acquired using a Zeiss LSM 710 confocal microscope. Images were subsequently processed using the ImageJ software package. For bacteria staining with filipin III, bacteria were cultured in LB medium (Lennox) overnight. Cultures were then centrifugated, resuspended in PBS with 50 µg/mL of filipin III and incubated for 3 hours at 37°C at 240 rpm.

### Live confocal microscopy

LAMP1-GFP HeLa were seeded into µ-Slide 8 wells glass bottom (80827 Ibidi) the day before experiment. The day of the experiment, cell medium was replaced with DMEM containing DND-99 rouge LysoTracker (L7528 fisher scientific) at 100 nM final concentration. Live experiments were performed on a Spinning disk confocal set up to incubate cells at 37°C with 5% CO_2_ during the experiment. Live imaging were performed for 1.5 hours with pictures taken every 15 minutes. Images were subsequently processed using the ImageJ software (47).

### Immunofluorescence labelling

DRG cells suspensions were plated on L-polylysin (Sigma)-coated 96-well plates. After 3 hours infection with OMVs, cells were fixed in PFA 4% and incubated at RT for 10 minutes. Cells were gently rinsed three times in PBS. LAMP1-GFP HeLa were seeded into 12 well plates with sterile slides in the bottom of the wells the day before the experiment. The day of the experiment, cells were treated as indicated in the legend of figures. After treatment, the cells were fixed with 4% PFA for 12 minutes at RT followed with 3 washes with PBS. The cells were then incubated 45 minutes at RT with MAXblock^TM^ Blocking Medium containing 0.3% Triton. Cells were then incubated with primary antibody against Galectin-1 (AB25138 abcam) or Galectin-3 (AB2785 abcam) at 1:500 and 1:100 respectively either 2 hours at RT or O/N at 4°C. The slides were then washed 3 times with cold PBS before being incubated with the Alexa Fluor secondary antibodies at 1:1000 for 1.5 hours at RT. After being washed 3 times with cold PBS the slides were mounted with Fluoroshield with DAPI (sigma F6057). Images were acquired using a Zeiss LSM 710 (Objective 100, CARL ZEISS SAS) or a Zeiss LSM 880 with an Airyscan module inverted confocal microscopes. Image analysis was performed using the ImageJ software and QuPath 0.6 software (47,48).

### Lipid extraction

We used a Folch-like extraction method to extract lipids from bacterial cells (49). Briefly, bacteria were grown under the same conditions as for OMVs production (see OMVs production above). After 8 hours culture at 37°C at 240 rpm shaking, the cultures were centrifuged at 6000 × g for 10 minutes to pellet the bacteria. The bacterial pellets were then resuspended in a 2:1 CHCl /MeOH solution and agitated for 1 hour at 40 rpm in a rotary shaker. Samples were then centrifugated for 10 minutes à 1 000 rpm to pellet bacterial cell debris. Supernatants containing lipids were washed with ultrapure water and agitated for 15 minutes at 40 rpm in a rotary shaker. Samples were then centrifuged for 10 minutes at 1 000 rpm and the organic phases were recovered and evaporated under N_2_. Dry lipids pellets were then resuspended in DMEM 1X without phenol red (21063 Gibco). As for OMVs, lipids concentration were quantified with a sulfo-phospho-vanillin reaction as described in Izard *et al* (46) with Triolein as a reference (44896-U Sigma Aldrich).

### Liposomes preparation

Liposomes were generated from mix of lipids derived from HlyF-producing bacteria (extracted as described above) and commercial lipids, namely 1-palmitoyl-2-oleoyl-sn-glycero-3-phosphoethanolamine (POPE) (850757P Avanti), 1-palmitoyl-2-oleoyl-sn-glycero- 3-phospho-(1’-rac-glycerol) (POPG) (840457P Avanti) and 1’,3’-bis[1,2-dipalmitoyl-sn-glycero-3-phospho]-glycerol (CL) (710333P Avanti) or 16:0-12:0 NBD PE (810154P Avanti Research) as described in Kehl *et al* (26). Before extrusion, lipidic mixes of 2 mg were prepared with 1 mg of lipids from HlyF-producing bacteria and 1 mg of lipids containing 60% of POPE, 30% of POPG and 10% of CL (26) resuspended each in chloroform/methanol (2/1, v/v). For liposomes-control, a lipidic mixes of 2 mg were prepared with commercial lipids containing 60% of POPE, 30% of POPG and 10% of CL. Lipidic mixes were then evaporated under N_2._ Dry lipids pellets were then resuspended in DMEM without phenol red (21063 Gibco). Liposomes were produced by extrusion, according to the manufacturer’s instruction about the Avanti Mini Extruder Extrusion Technique (610000-1EA Avanti Polar Lipids Inc.). Lipids mixes were extruded using 200 nm (610006-1EA Avanti) then 100 nm membranes (610005-1EA Avanti). When indicated, liposomes were then labelled with 1 µL of DiI (V22885 ThermoFisher) for 100 µL of liposomes suspension at 37°C during 30 minutes and then stored at 4°C. Images were acquired using a Zeiss LSM 880 confocal microscope with an Airyscan module. Images were subsequently processed using the ImageJ software (47).

### Statistical analysis

All graphical figures and statistical analyses were performed with the GraphPad Prism 10 software (GraphPad Software, San Diego, CA).

## Data availability

No reagents, plasmids, bacterial strains or cell models were generated for this study. All the materials used in this study are available from the lead contact with a completed materials transfer agreement.

## Acknowledgements

This project was supported by funding from the French National Research Agency projects SMERSEC (ANR-20-CE18-0016) and VeSPath (ANR-23-CE14-0070). This project also benefited from a doctoral fellowship provided by the French Ministry of Higher Education and Research. We are indebted to Pr. Andrea Ballabio for kindly providing the LAMP1-GFP HeLa cell line. Imaging experiments were performed at the Infinity-INSERM UMR1291 core facility connected to Toulouse Réseau Imagerie network, member of the France-BioImaging national infrastructure supported by the French National Research Agency (ANR-24-INBS- 0005 FBI BIOGEN). We thank Sophie Allart from the imaging platform of Infinity. We gratefully acknowledge Thomas Mangeat and Brice Ronsin from the light imaging platform of the CBI, and Justine Creff for their precious help with pseudo high resolution confocal microscopy.

## Author contributions

Conceptualization: L.D., E.O.; methodology: P.B., L.D., E.O., C.P., F.T. ; investigation: C.B., L.D., L.L, S.L., C.M., C.P., C.R. ; funding acquisition: E.O., L.D. ; writing original draft: L.D., E.O., C.P.

## Declaration of interests

The authors declare no competing interests

## Supplemental information

Document S1. Supplemental figures S1-4 :

Figure S1: Treatment with OMVs or lipids from HlyF-producing bacteria induce a time-dependent loss of acidity in lysosomes

Figure S2: Filipin III does not stain bacterial lipids.

Figure S3: OMVs from HlyF-producing bacteria specifically induce lysosomal recruitment of galectin-3

Figure S4: Fluorescent lipids from HlyF-producing bacteria are integrated in lysosomes membranes.

## Declaration of generative AI and AI-assisted technologies in the writing process

During the preparation of this work, the author(s) used DeepL and DeepL write for improving the English wording of the text. The author(s) reviewed and edited the output as needed and take full responsibility for the content of the published article.

## Notes

### Competing Interest Statement

The authors have declared no competing interest.

