## Supplementary figures for "Bacterial lipid effectors subvert host autophagic flux by inducing lysosomal dysfunction"

Supplementary figures S1-4

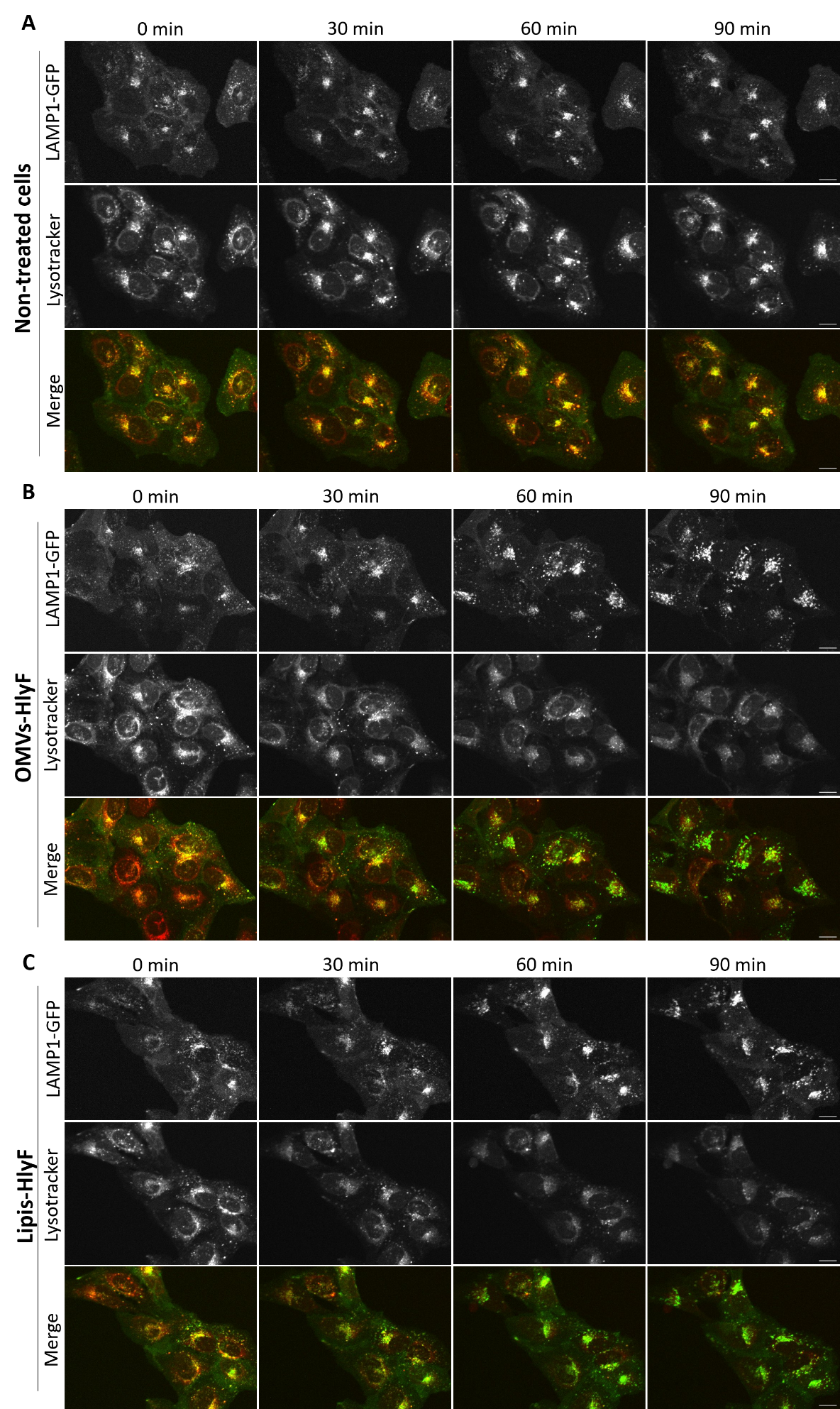

**Figure S1: Treatment with OMVs or lipids from HlyF-producing bacteria induce a time-dependent loss of acidity in lysosomes.**

**A-C** HeLa LAMP1-GFP cells were (**A**) left untreated, (**B**) treated with OMVs-HlyF at 10 µg/mL of lipids or (**C**) with lipids-HlyF at 10 µg/mL. LAMP1-GFP and LysoTracker signals are displayed in green and red respectively in the merged channel. Experiments were performed with a Spinning disk confocal set up to incubate cells at 37°C with 5% CO_2_ during the experiment and pictures were taken at the beginning of the experiment (0), 30min, 60min and 90min. At least 50 cells were analyzed per condition. Images representative of at least 3 independent experiments. Scale bar = 16 µm.

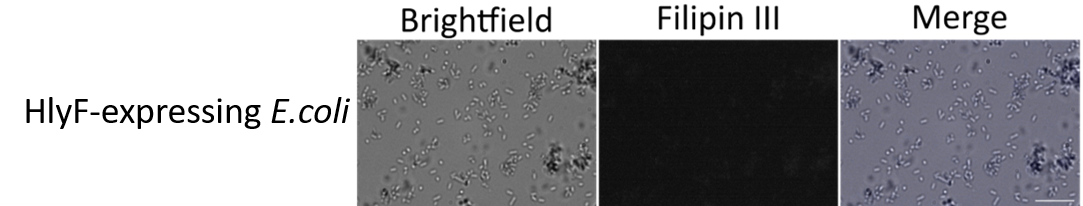

**Figure S2: Filipin III does not stain bacterial lipids.** Bacteria expressing HlyF from the BL21 strain were fixed to a poly-lysine-coated glass slide before being labelled with 50 µg/mL of filipin III. Scale bar = 10µm.

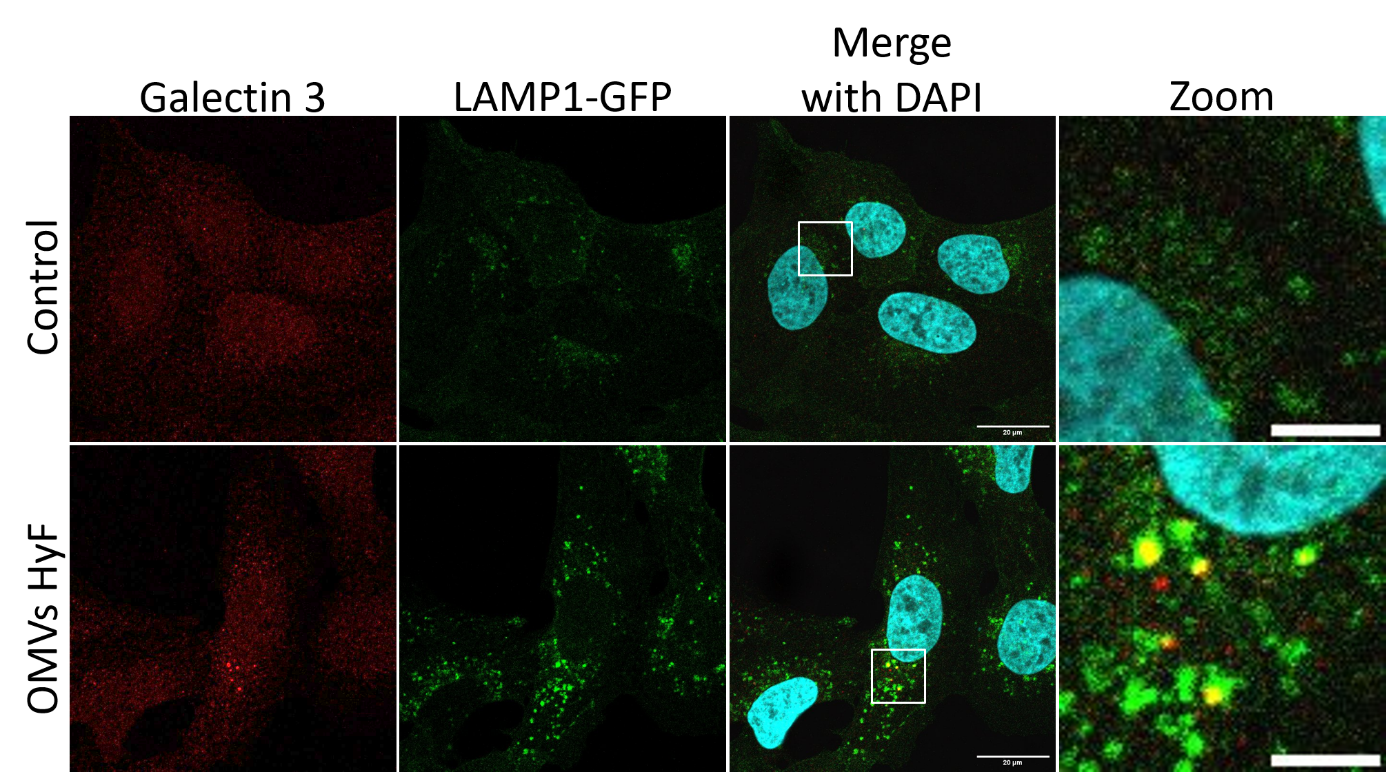

**Figure S3: OMVs from HlyF-producing bacteria specifically induce lysosomal recruitment of galectin-3**

HeLa LAMP1-GFP cells were treated for 1.5 hour with OMVs-HlyF at 10 µg/mL lipids or left untreated. Cells were then fixed and labeled with an anti-Galectin-3 antibody. Galectin-3 and LAMP1-GFP are displayed in red and green respectively in the merged channel. A total of 50 cells were analyzed per condition. Images are representative of three independent experiments. Scale bar = 20 µm. Boxed regions are shown at higher magnification (scale bar = 5µm).

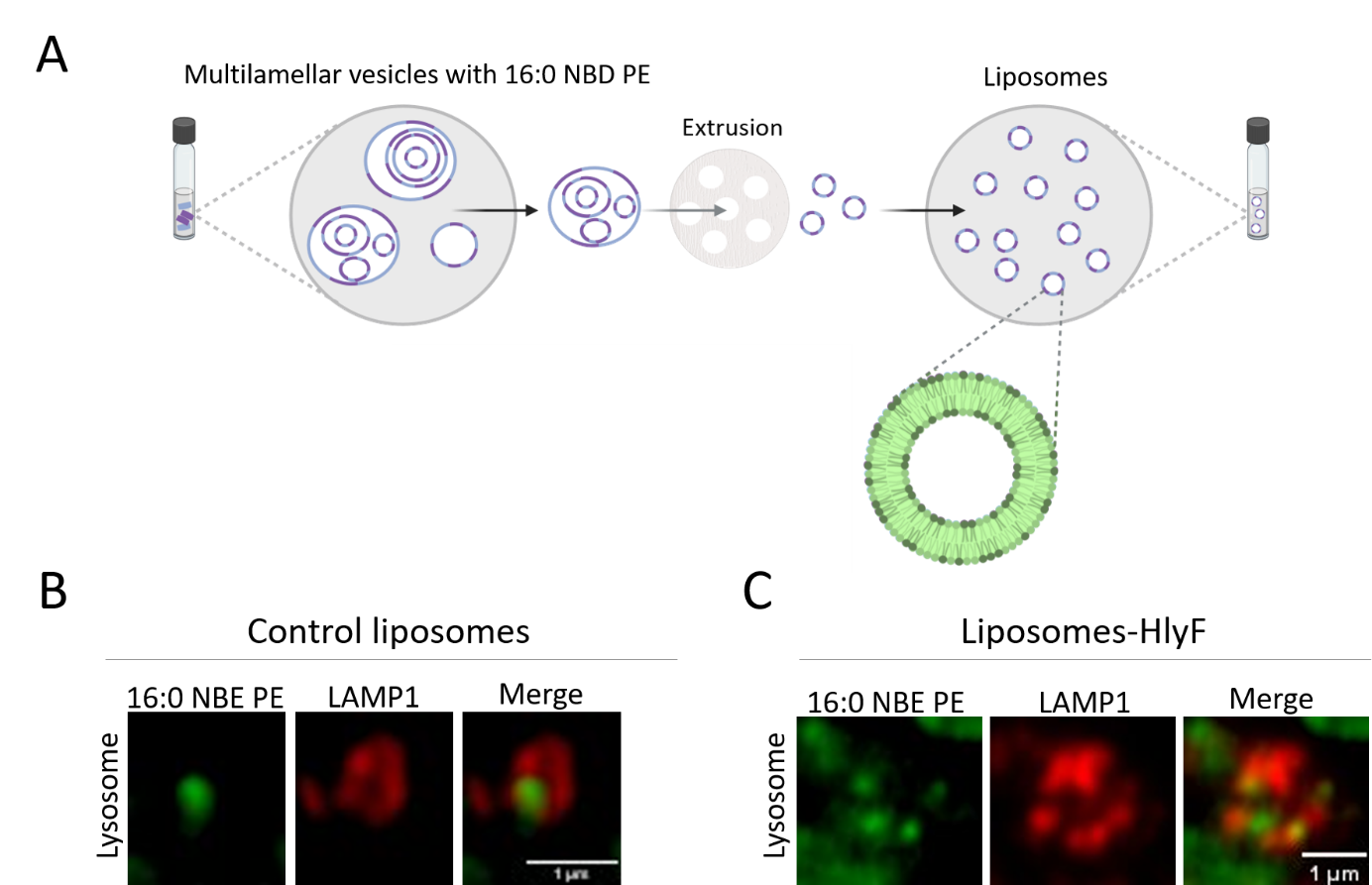

**Figure S4: Fluorescent** **lipids from HlyF-producing bacteria are integrated in lysosomes membranes.**

**A**. To prepare liposomes, lipids-HlyF and the fluorescent lipid 16:0 NBD PE were resuspended in aqueous media and then extruded through 200 then 100 nm filters. After extrusion, liposomes present a normalized size. **B**. HeLa LAMP1-GFP were treated with 100µg/mL of fluorescent liposomes made with 50% of synthetic lipids and 50% of lipids-HlyF (liposomes-HlyF) or 50% of lipids from control bacteria (control liposomes) during 3 hours. Images were taken with a Zeiss 880 confocal microscope with an Airyscan module. Scale bar = 1 µm.
